# Development and validation of a high throughput screening platform to enable target identification in skeletal muscle cells from Duchenne Muscular Dystrophy (DMD) patients

**DOI:** 10.1101/2023.05.24.542079

**Authors:** Santosh Hariharan, Oana Lorintiu, Chia-Chin Lee, Eve Duchemin-Pelletier, Xianfeng Li, Aileen Healy, Regis Doyonnas, Luc Selig, Pauline Poydenot, Erwann Ventre, Andrea Weston, Jane Owens, Nicolas Christoforou

## Abstract

Duchenne muscular dystrophy (DMD) is a progressive and fatal muscle degenerating disease caused by dystrophin deficiency. Effective methods for drug discovery for the treatment of DMD requires systems to be physiologically relevant, scalable, and effective. To this end, the Myoscreen platform offers a scalable and physiologically relevant system for generating and characterizing patient-derived myotubes. Morphological profiling is a powerful technique involving the simultaneous measurement of hundreds of morphological parameters from fluorescence microscopy images and using machine learning to predict cellular activity. Here, we describe combining the Myoscreen platform and high dimensional morphological profiling to accurately predict a phenotype associated with the lack of Dystrophin expression in patient derived myotubes. Using this methodology, we evaluated a series of Dystrophin-associated protein complex (DAPC) candidates and identified that the combination of Utrophin and α- Sarcoglycan yielded highest morphological differences between DMD and non-DMD donors. Finally, we validated this methodology by knocking down Dystrophin expression in non-DMD cells as well as introducing Dystrophin expression in DMD cells. Knocking down Dystrophin in non-DMD cells shifted their morphological profile to one that is similar to DMD cells while introducing Dystrophin in DMD cells shifted their morphological profile towards non-DMD cells. In conclusion, we have developed a platform that accurately predicts the DMD disease phenotype in a disease relevant cell type. Ultimately this platform may have wide applications in the drug development process include identification of disease modifier genes, screening of novel therapeutic moieties, and as a potency assay for future therapeutics.

## INTRODUCTION

Duchenne muscular dystrophy (DMD) is a genetic disease characterized by progressive muscle weakness and degeneration (Verhaart & Aartsma-Rus, 2019). It is caused by mutations in *DMD*, a gene located on the X chromosome, spanning a genomic range of greater than 2 Mb, and encoding for dystrophin, a component of the dystrophin-associated protein complex (DAPC), which bridges the inner cytoskeleton and the extracellular matrix; this is a disease that primarily affects boys (Dowling et al., 2021). Most pathogenic mutations result in a loss of mRNA stability or the introduction of a premature codon, ultimately resulting in the lack of Dystrophin.

The DAPC is a multi-component protein complex that resides within the cell membrane, or sarcolemma, of the myofiber (Campbell & Kahl, 1989; Ervasti et al., 1990). The DAPC has three main functions: (i) it shields the sarcolemma from stresses that occur during muscle contraction, (ii) it links the intracellular cytoskeletal scaffold to the extracellular matrix and participates in the transmission of force between the two compartments, and (iii) it functions as a scaffold for signaling molecules (Dowling et al., 2021). It is comprised of 11 main protein components including Dystrophin, α-Dystroglycan, β-Dystroglycan, α-Sarcoglycan, β-Sarcoglycan, γ- Sarcoglycan, δ-Sarcoglycan, Sarcospan, Syntrophin, Dystrobrevin, and Nitric Oxide Synthase (Duan et al., 2021).

Dystrophin interacts with the DAPC and the sarcolemma mainly through one of the DAPC components, β-Dystroglycan (Ervasti & Campbell, 1991) and this interaction stabilizes the complex and the sarcolemma and enables its proper function. Loss of Dystrophin due to pathogenic mutations results in significant changes in the expression and localization of both scaffolding and signaling proteins associated with the DAPC (Allen et al., 2016); DAPC disassembly and loss of the interactions between the intracellular scaffold (F-Actin) and the extracellular matrix (Duan et al., 2021). Ultimately this results in sarcolemma tearing upon muscle contraction, development of abnormal Ca^2+^ homeostasis, long-term deficits in muscle regeneration, and tissue fibrosis (Burr & Molkentin, 2015; Chan & Head, 2011; Petrof et al., 1993; Schmalbruch, 1984).

Understanding the disease process and ultimately identifying targets that may function as modifiers within this process requires models that accurately capture key pathophysiological processes at the cellular level. Most of the research and discovery related to DMD has historically been pursued in the *mdx* mouse, an animal with a spontaneous dystrophin mutation (Gerde & Scholander, 1987; Partridge, 2013). A significant limitation of the *mdx* model is that it presents with less severe disease process and does not closely recapitulate the human phenotype (Partridge, 2013). The development of improved preclinical models that carry human disease-relevant mutations which are based on the formation of differentiated and functional human myofibers is key for enabling the discovery process. Additionally, it is essential that readouts developed within such a platform are reproducible in the context of a range of pathogenic mutations and allow for a robust assay window to reduce noise and enable the discovery of new targets with high confidence.

For discovery of high confidence targets for muscular dystrophy, it is important to develop physiologically relevant disease models that can accurately mimic in-vivo functionality. While 2D cell culture models are scalable, they do not recapitulate the myofiber morphology and functionality, making the system less valuable for target and drug discovery. Engineered micropatterned plates can provide the appropriate physical environment for myogenesis (Li et al., 2008). Skeletal muscle cells differentiated in micropatterned plates form mature myotubes that are aligned and striated and express dystrophin (Young et al., 2018). Therefore, micropatterned plates are ideal for target and drug screening owing to their physiological relevance for myotube formation and by enabling imaging with quantification of complex endpoints. Image based single cell morphological profiling has gained popularity over the past few years. With automated microscopy, a large set of images can be acquired in a very short time. Morphological profiling involves computing hundreds of single cell image features and using complex machine learning and/or deep learning models to uncover novel biology (Carpenter et al., 2006; Dao et al., 2016). The high dimensional morphological profile can reveal enhanced information about the cellular perturbation compared to the biased single readout from images. Morphological profiling has been used in the past to identify the mechanism of action of test compounds as well as disease mechanisms (Scheeder et al., 2018; Ziegler et al., 2021). Recently, unbiased morphological profiling has also been used for identifying signatures of disease phenotypes (Schiff et al., 2022) as well as to distinguish healthy and diseased populations (Oppermann et al., 2016; Wali et al., 2021).

Here we describe the development and validation of a platform based on the implementation of primary human DMD patient and control (non-DMD) myoblasts which are induced to differentiate within a micropatterned substrate allowing for the highly reproducible formation of differentiated myotubes. Furthermore, we demonstrate that by morphological profiling in differentiated myotubes, immunostained for Utrophin and α-Sarcoglycan, we can consistently distinguish the two populations (DMD vs. non-DMD) and provide evidence that this is attributed to the absence of dystrophin.

## RESULTS

### Characterizing primary donor cells for MyoScreen platform

We evaluated a set of human derived skeletal myoblasts for further studies based on three success criteria: (1) donors were selected based on sufficient availability of starting material with a minimum number of cryopreserved vials containing early passage (P0) myoblasts being set at n≥3. This is to ensure that a sufficient supply of cells is available for the entire experiment and to limit batch-to-batch variability, (2) donors were selected based on being male to ensure disease relevance phenotypes observed in DMD as the disease primarily affects boys and mitigate any effect on morphology based on sex, and (3) donors were selected based on their age (3-5 years) since the disease affects boys at a similar age. Patient donor cells that met the 3 criteria are included in Table 1. Further triaging was pursued for donors that met the three criteria with the goal of selecting 2 DMD and 2 non-DMD donors using a fusion index (> 0.1) for DMD donors and expression of Dystrophin in non-DMD donors.

**Table 1.** Donor selection metrics for different donors that were evaluated for the study. . Donor selection metrics for different donors that were evaluated for the study.

| Donor | Age | Diagnosis | Mutation | Received Material | Freezing (MCB) | Master Cell Bank Characterization |  | Freezing (WCB) | Working Cell Bank Characterization |  |
| --- | --- | --- | --- | --- | --- | --- | --- | --- | --- | --- |
|  |  |  |  | Desmin Pos (%) | Desmin Pos (%) | Desmin Pos (%) | Fusion Index | Desmin Pos (%) | Desmin Pos (%) | Fusion Index |
| DMD #1 | < 3 yrs | DMD | Deletion, Exons 18-44 | 73 | 85 | 75 | 0.35 | 70 | 65 | 0.33 |
| DMD #2 | 3 - 6 yrs | DMD | Mutation c.5602_5605del, Asp1866Glufs*5 | 65 | 80 | 89 | 0.46 | 89 | 88 | 0.37 |
| DMD #3 | 3 - 6 yrs | DMD | Mutation c.4729C>T, p.Arg1577* | 69 | 83 | 73 | 0.14 | 77 | 67 | 0.17 |
| DMD #4 | < 3 yrs | DMD | Deletion, Exons 46-52 | >80 | >80 | >80 | >0.3 | >80 | 94 | 0.5 |
| DMD #5 | < 3 yrs | DMD | Deletion, Exons 46-55 | 76 | 53 | 30 | 0.2 | 27 | 16 | 0.11 |
| DMD #6 | 3 - 6 yrs | DMD | Hemizygote Duplication Exons 61-68 | 77 | 50 | 37 | 0.003 | 26 | N/A | N/A |
| DMD #7 | 3 - 6 yrs | DMD | Deletion, Exons 6-7 | 70 | 68 | 40 | 0.17 | 29 | 14 | 0.02 |
| DMD #8 | 3 - 6 yrs | DMD | N/A | 75 | 16 | 3 | 0.002 | 1 | N/A | N/A |
| DMD #9 | 3 - 6 yrs | DMD | Mutation c.7729G>T, p.Glu2577* | 63 | 53 | 23 | 0.19 | 20 | 12 | 0.03 |
| Non-DMD #1 | < 3 yrs | Non-DMD | N/A | 95 | 98 | 99 | 0.52 | 99 | 98 | 0.43 |
| Non-DMD #2 | 6 - 9 yrs | Non-DMD | N/A | 74 | 85 | 88 | 0.44 | 84 | 82 | 0.35 |
| Non-DMD #3 | 6 - 9 yrs | Non-DMD | N/A | 89 | 93 | 98 | 0.56 | 96 | 97 | 0.7 |
| Non-DMD #4 | 3 - 6 yrs | Non-DMD | N/A | 89 | 42 | 37 | 0.009 | 30 | N/A | N/A |

The process of donor-specific myoblast selection and cell expansion is illustrated in Figure 1A. Briefly, myoblasts were isolated at a hospital setting from patients following the collection of a skeletal muscle biopsy. The cells were subsequently sorted using magnetic beads based on expression of CD56. CD56 (Neural Cell Adhesion Molecule – NCAM), is a glycoprotein expressed on the surface of skeletal muscle cells (Walsh & Ritter, 1981). Cell sorting was then implemented to quantify the proportion of myoblasts by quantifying Desmin positive cells (DES^pos^). Cell populations that were at least 25% DES^pos^ were selected for further expansion. Patient cells obtained at the hospital were expanded and cryopreserved into master and working banks. Before creating a master or a working bank, cells were again characterized for Desmin expression. This was done to ensure that the freeze-thaw process did not alter the proportion of DES^pos^ cells in the cryopreserved cell population. Cells from the working bank were subsequently used for all downstream experiments. In all, cells from 4 donors were selected with 2 coming from DMD patients and the other 2 coming from non-DMD individuals.

**Figure 1.**
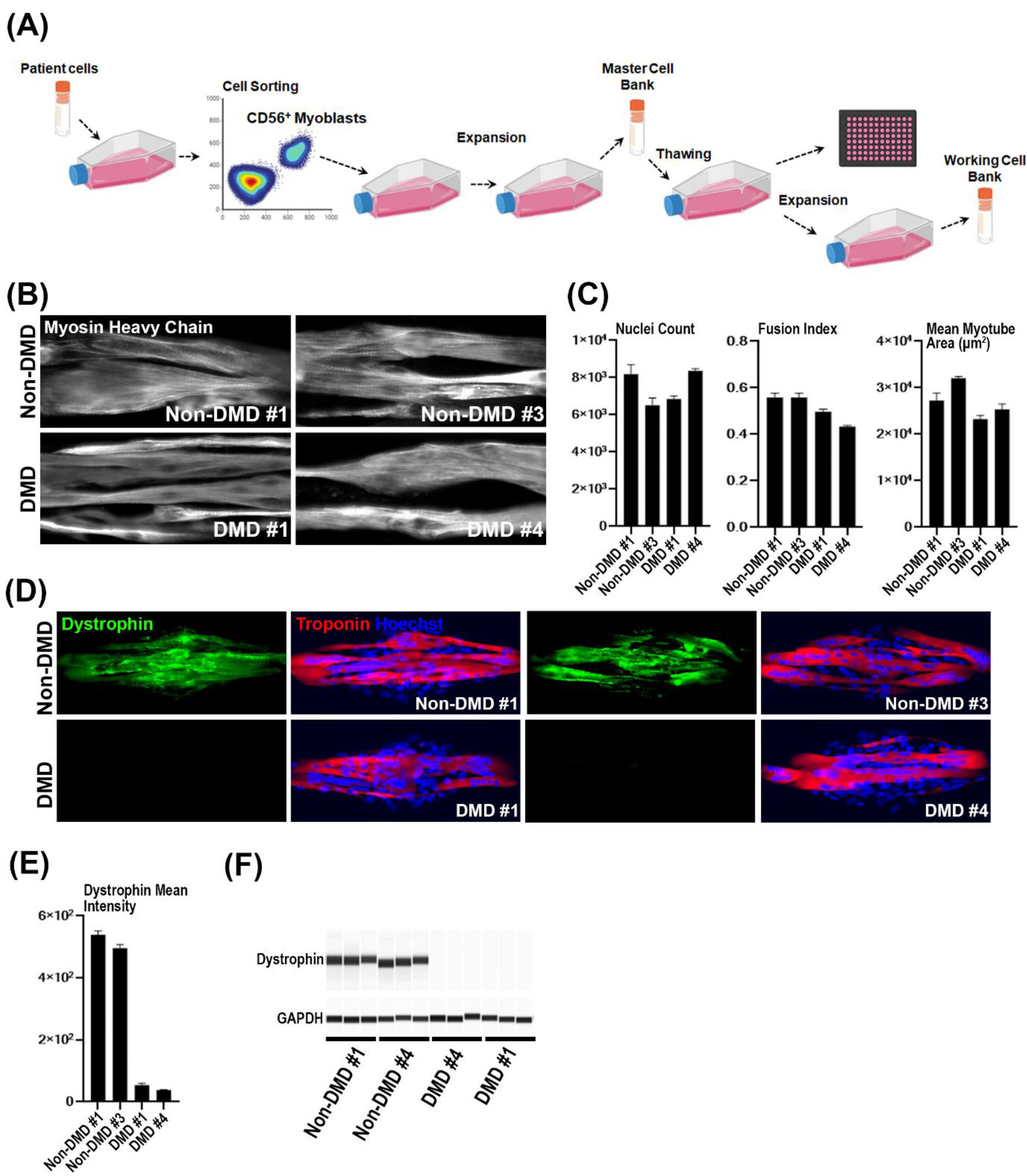
Donor selection, myotube quantification and dystrophin expression validation. **(A)** Primary myoblasts from DMD and non-DMD donors were selected among other donors based on retaining their capacity to differentiate (% Desmin^+^ myoblasts and Fusion Index) into myotubes within the MyoScreen platform. Cells expanded following patient biopsy collection were subsequently enriched for myoblasts using flow cytometry (CD56^+^). Primary vials were sourced, thawed, and the proportion of Desmin^+^ cells was determined using flow cytometry. Cells were expanded and cryopreserved into master banks (MB) at which point they were characterized using flow cytometry (Desmin^+^ cells) and the MyoScreen platform (fusion index). Finally, master bank vials were thawed, expanded, and finally cryopreserved into working cell banks (WB) at which point they were characterized using flow cytometry (Desmin^+^ cells) and the MyoScreen platform (fusion index). Cells were selected based on consistency in their doubling time, proportion of Desmin^+^ cells and fusion index. **(B)** Myoblasts from the two non-DMD (Non- DMD #1, Non-DMD #3) and two DMD (DMD #1, DMD #4) donors were differentiated for 10 days in MyoScreen micropatterned plates. The differentiated myotubes were stained for Myosin Heavy Chain (MHC) and Dystrophin protein. Sample images of individual Myoscreen islands for each donor stained for Myosin heavy chain (MHC). **(C)** Quantification of nuclei count, myotube area and fusion index for myotubes generated from the 4 different donors. Overall, the fusion index which quantifies muscle cell myotube formation, is significantly lower in DMD donor cells compared to non-DMD donors. **(D)** Sample fluorescent images from the 4 donors stained for dystrophin. Dystrophin was not observed with this endpoint in differentiated myotubes from DMD donors. **(E)** Quantification of dystrophin intensity from DMD and non-DMD donors. **(F)** Dystrophin protein expression was further evaluated using Western blotting.

For further characterization, the cells from the master bank were differentiated and stained for Myosin heavy chain (MHC) to identify myotubes (Figure 1B). Cell images were then used to extract the myotube area (MHC-positive area), and nuclei count (Figure 1C), which was used to compute the fusion index (See Methods). As expected, the fusion index for the DMD donors (DMD #1 and DMD #4) was significantly lower compared to non-DMD donors (Non-DMD #1 and Non-DMD #3) (Meng et al., 2020).

Dystrophin expression was characterized by immunofluorescence (IF) and simple western blot (Figures 1D-1F). Nuclei and myotubes were identified using Hoechst and Troponin T respectively. The DMD donors have very low to no expression of Dystrophin compared to the non-DMD donors.

### Image based profiling of donor myotubes separates the donor population better than protein expression

Next, we sought to identify specific proteins that were differentially expressed between DMD and non-DMD myotubes. To this end, we screened for proteins associated with the Dystrophin Associated Protein Complex (DAPC). Myoblasts were seeded and differentiated within the Cytoo MyoScreen 96-well micropatterned plates (Bosch-Fortea et al., 2019; Young et al., 2018). Selecting the platform allowed us to (i) implement a physiologically relevant system for myotube formation and (ii) achieve consistency in the differentiation of myotubes between the various donors.

To examine for potential differences between DMD and non-DMD myotubes we compared protein expression levels for several DAPC proteins using immunofluorescence (Figure S2.1). Immunofluorescence staining against Troponin T was used to identify differentiated myotubes within individual micropatterns (Figure 1D). Cells were counterstained for the various DAPC components. Using image analysis, we determined the average pixel intensity for each myotube. The intensity was then computed for myotubes across each micropattern and each well. The sample average and standard deviation for each donor and DAPC protein were determined (Figure S2.1). In the case of Utrophin, donor DMD #4 had significantly (p-value = 0.0021 (Non-DMD #1); p-value = 0.0001 (Non-DMD #3) higher Utrophin intensity compared to non-DMD donors (Non-DMD #1 and Non-DMD #3). However, this was not true for the DMD donor DMD #1. The variability in expression levels may be attributed to the differences in the severity of DMD in the donor cells (Janghra et al., 2016). We observed similar variability in expression levels for the other DAPC proteins (Figure S2.1). Taken together, the expression levels of DAPC proteins did not display significant changes between the DMD and non-DMD donor population.

We speculated that even though we did not detect significant differences in DAPC protein expression levels using immunofluorescence, there may exist significant differences in the manner via which these proteins organize between the two cell populations (DMD vs. non-DMD). To this end, we implemented cell image-based profiling to further analyze the captured images. To quantify the ability of individual DAPC proteins to separate the two populations, we stained for each DAPC marker separately along with Troponin T and Hoechst stain on the MyoScreen platform. CellProfiler was used for segmentation of myotubes and extracting 235 features from individual myotubes pertaining to cell pixel intensity, pixel intensity distribution, texture, and granularity (See Methods). The extracted features were pre-processed and normalized. To avoid the effect of nuclear or Troponin T channel morphology, we restricted ourselves to features from the respective DAPC protein image.

To visualize the high dimensional data, we used t-SNE algorithm (van der Maaten & Hinton, 2008). T-SNE is an embedding method which is designed to preserve the high dimensional intra- point distances in a low dimensional (2D) space. The t-SNE plots are shown in Figure 2A and supplemental Figure S2.2. Each t-SNE plot is derived from features based upon specific DAPC markers. The t-SNE plot reflects the ability for the individual DAPC markers to separate the donor cells of DMD (Red) and non-DMD (Green) populations. DAPC protein Utrophin and α-Sarcoglycan demonstrated improved separation between the two populations compared to the other DAPC markers (Figure 2A and Supplemental Figure S2.2). Amongst all the DAPC proteins that were evaluated, Utrophin achieved the highest degree of separation visually.

**Figure 2.**
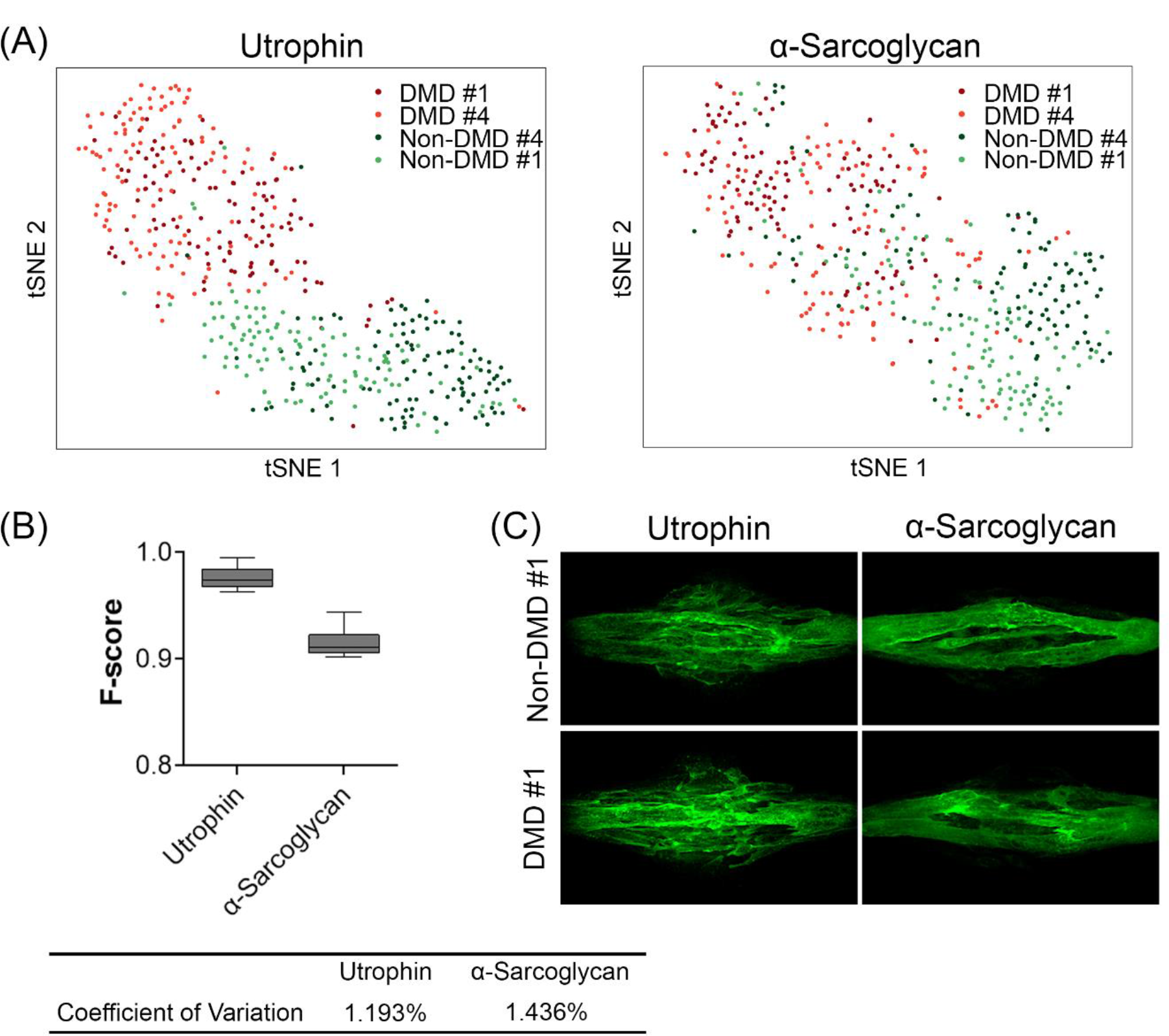
**Cell profiling of myotubes using only the individual (Utrophin and α-Sarcoglycan) DAPC marker features**. **(A)** t-SNE plots of multi-dimensional cell profiler features. The left panel is derived using only the utrophin channel features while the right panel is derived using the α- Sarcoglycan features. Each point is a single myotube and the color indicates the donors. Utrophin displays a higher separation on the t-SNE plot compared to α-Sarcoglycan. **(B)** Support vector Machine (SVM) 10-fold cross validation data confirms higher separation with Utrophin features compared to α-Sarcoglycan. Further the coefficient of variation (CV) is lower for Utrophin. **(C)** Sample images of fluorescent labelled images for DAPC proteins utrophin and α-Sarcoglycan for Non-DMD #1 (non-DMD) and DMD #1 (DMD) donors. There was no difference between average intensity of utrophin and α-Sarcoglycan between the DMD and non-DMD donors (Supplementary Fig S2.1).

To quantify the separation, we entered the DAPC image features to an SVM classifier and performed 10-fold cross validation. For performing cross-validation, the data was divided into 10 parts. 9 parts were used for training and 1 part was used for testing the accuracy of the model. The process was repeated 10 times with different parts of the data being used for either testing or training. Robust performance was determined by higher accuracy and lower dispersion in accuracy among the cross validation runs. The accuracy was computed via calculating the F-score (See methods). The F-score for Utrophin and α-Sarcoglycan is shown in Figure 2B. We determined that the F-score for Utrophin was significantly higher when compared to that of α-Sarcoglycan. The F-score for other DAPC markers was lower when compared to that of Utrophin and α- Sarcoglycan.

We wanted to further evaluate whether the separation observed when using Utrophin and α- Sarcoglycan is independent of the MyoScreen platform. We plated DMD and non-DMD donor cells in a standard flat-bottom 96 microwell plate and used the morphological profiling to compare the result to cells plated in the MyoScreen platform. We observed that the MyoScreen platform combined with morphological profiling had a significantly higher F-score compared to cell plated in regular 96 microwell plates (Supplemental figure S2.3).

Taken together, our results indicate that use of morphological profiling in combination with the MyoScreen platform can yield higher separation between the DMD and non-DMD populations while using the DAPC proteins as a readout.

### Combining DAPC markers yields better discrimination between DMD and non-DMD populations

DAPC proteins interact with each other on the cell membrane to ensure proper functioning in myotubes. In healthy myotubes, dystrophin interacts with actin (N-terminus) and the DAPC complex (C-terminus), which in turn interacts with laminin proteins in the extracellular space. The unique interaction of dystrophin with DAPC proteins renders structural integrity to myotubes (Bhat et al., 2018; Gao & McNally, 2015). Previously we examined individual DAPC proteins using morphological profiling. Here, we hypothesized that capturing these DAPC protein interactions would yield significantly better separation between DMD and non-DMD populations compared to individual proteins. This is based on the premise that it is the loss of dystrophin that disrupts the interaction of the multi-faceted DAPC proteins (Duan et al., 2021). Quantifying the changes in the interaction could yield significantly improved separation between the populations.

Instead of pursuing all possible combinations, we restricted ourselves to testing two DAPC protein components at a time. Performing experiments with all possible interactions of two markers would result in 21 combinations. To prioritize DAPC combinations, we approached the problem in a data-driven manner. We first ranked the DAPC markers that demonstrate the highest degree of separation between the two populations. From that ranking we selected the top two markers against which we tested all other protein combinations. This resulted in Utrophin and α-Sarcoglycan as the two proteins with the highest F-score and low coefficient of variation. We evaluated the remaining DAPC proteins (Syntrophin, δ-Sarcoglycan, α- Dystroglycan, β-Dystroglycan) in combination with Utrophin or α-Sarcoglycan. For each of the combinations, we computed F-scores across 10-fold cross validation using individual markers alone as well as combined DAPC markers. As before, the F-score was computed by pooling the DMD donors as one class and the pooled non-DMD donors as another class. The F-scores for the combination of markers with utrophin is shown in Figure 3A. For almost all combinations, the combined marker dataset was better at separating the DMD and non-DMD population compared to an individual marker alone. Combining Utrophin and α-Sarcoglycan yielded the highest separation (F-Score .99). This is not surprising as Utrophin and α-Sarcoglycan individually were able to separate the two populations with greater than 0.9 F-score. Combination of other markers (δ-Sarcoglycan, α-Dystroglycan and β-Sarcoglycan) with Utrophin also yielded higher separation between the two populations, even though δ-Sarcoglycan, α-Dystroglycan and β- Sarcoglycan performed very poorly in separating the two populations individually. The t-SNE plot for the data from marker combination is shown in Figure 3B. We used shades of red to denote the DMD donor population while shades of green to denote the non-DMD donor population. As shown from the t-SNE plots, there is a clear separation between the red (DMD) and green (non-DMD) populations when using Utrophin as constant for the combination of DAPC protein. We also performed combination analysis with α-Sarcoglycan along with α-Dystroglycan, β- Sarcoglycan and δ-Sarcoglycan (Figures 3C-D). Unlike Utrophin, the combination of DAPC proteins with α-Sarcoglycan did not always yield significantly higher F-scores compared to the individual DAPC markers alone (compare red, blue and green boxes), implying that most of the combination effect is driven by α-Sarcoglycan. The median F-score for the combination with α- Sarcoglycan was also lower when compared to combination of DAPC proteins with Utrophin. In addition, we also tested combinations of δ-Sarcoglycan and α-Dystroglycan which also resulted in lower F-scores.

**Figure 3.**
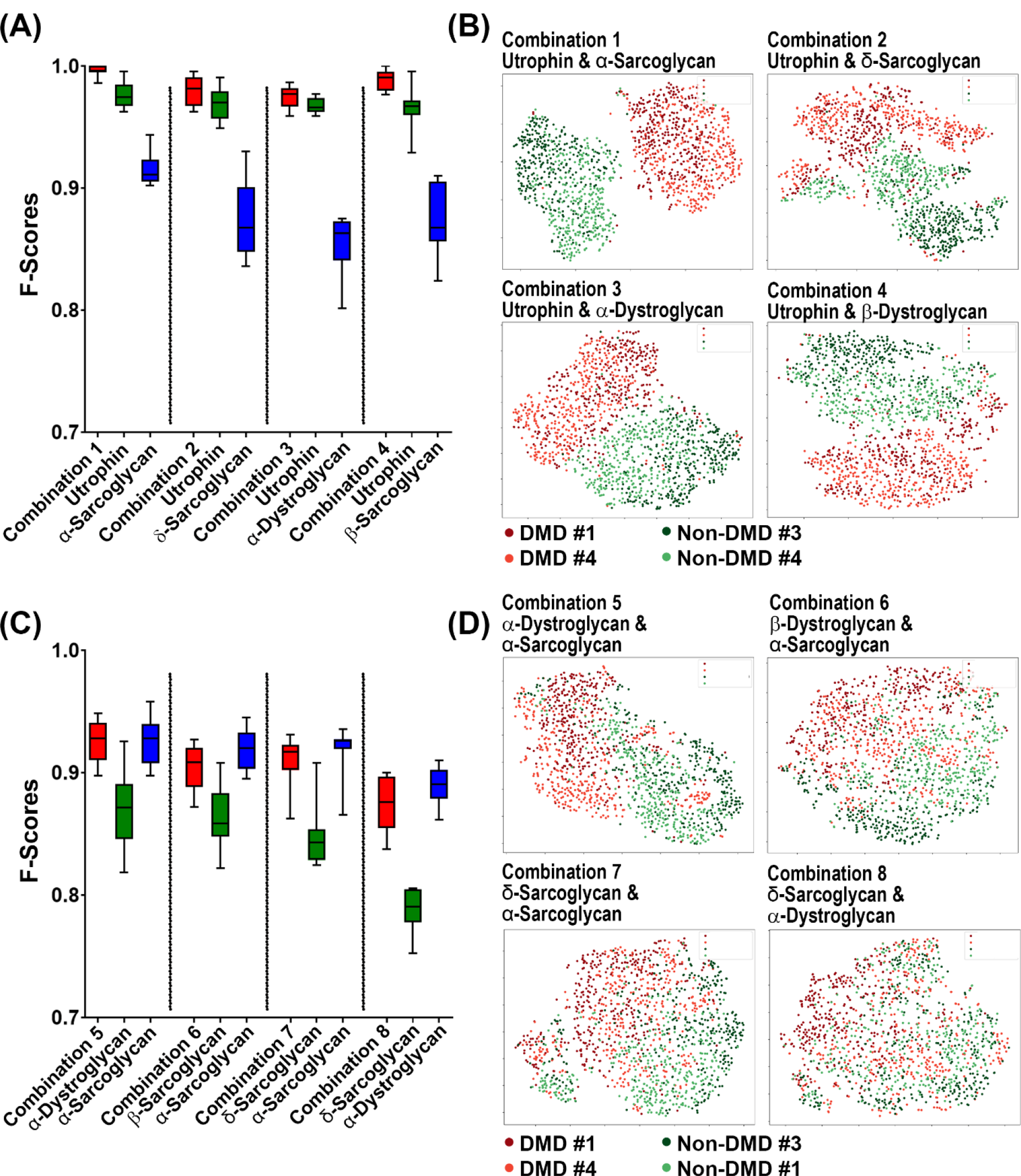
**DAPC protein combinations display superior performance when used in combination with Utrophin**. **(A)** Cross validation F-Score from morphological features of other DAPC proteins in combination with utrophin from cells cultured on MyoScreen platform. Combinations of DAPC proteins with Utrophin achieve better performance in separating the DMD and the non-DMD donor population when compared to individual DAPC proteins. Utrophin in combination with α- Sarcoglycan displayed much higher performance (F-Score .99) compared to either single DAPC protein alone or other DAPC protein combinations with Utrophin. **(B)** t-SNE plots showing donor cells from the MyoScreen platform for DAPC protein combinations. Comparing the t-SNE plots, the high separation between the DMD and non-DMD cells is highly evident in “Combination-1” (Utrophin + a-Sarcoglycan) compared to other DAPC protein combinations with utrophin. **(C)** Cross validation using the subcellular features using DAPC proteins in combination with α- Sarcoglycan obtained from cell profiling from cells cultured on the MyoScreen platform. Combinations of DAPC proteins with α-Sarcoglycan also shows better performance in separating the DMD and the non-DMD donor population when compared to individual DAPC alone. **(D)** t-SNE plots showing donor cells from the MyoScreen platform for DAPC protein combinations.

The combinations of DAPC proteins with Utrophin have higher F-scores compared to combinations of DAPC proteins with α-Sarcoglycan. For α-Sarcoglycan, the combination of DAPC proteins yielded similar F-scores compared to α-Sarcoglycan alone, except for combinations of α-Dystroglycan and δ-Sarcoglycan. Taken together, we determined that combining two DAPC markers yielded in a significantly better separation of DMD and non-DMD populations.

### siRNA-mediated knock-down of Dystrophin in healthy donors shifts to a DMD like profile

We hypothesized that the differences detected when comparing DMD and non-DMD myotubes could be attributed to the lack of Dystrophin. Knocking down Dystrophin in non-DMD cells could normalize the differences between the two cell populations. To test this hypothesis, we transfected differentiating non-DMD cells with siRNAs targeting human Dystrophin and achieved a robust decrease (> 75% knock down) in Dystrophin expression (Figure 4A).

**Figure 4.**
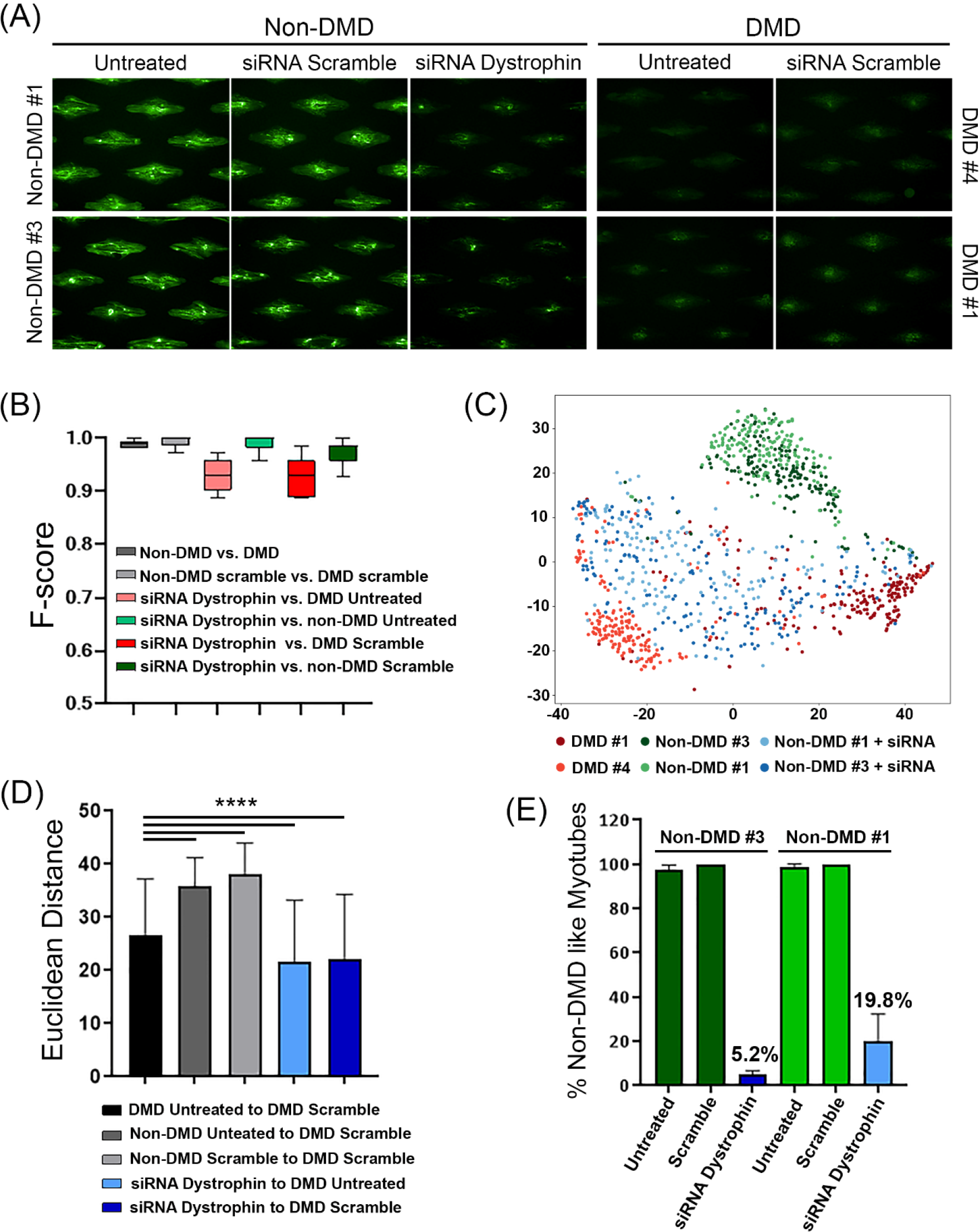
**Quantification of functional relevance of Dystrophin knock down (KD) using cell profiling**. **(A)** siRNA-based knock down of Dystrophin in non-DMD differentiated myotube cells (Non-DMD #1 and Non-DMD #3). The image intensity reflects dystrophin expression. For comparison, Dystrophin expression from the DMD-donors is included on the right. This is a baseline level of Dystrophin expression. In comparison, we saw that Dystrophin expression using siRNA3 is comparable to Dystrophin expression detected from the DMD donors. All images are pseudo colored. **(B)** F-score from cross validation analysis using SVM linear kernel by taking the 2 different categories at a time using Utrophin and α-Sarcoglycan features. As evident from the F-score, we can separate siRNA3 samples from both DMD and non-DMD scramble siRNA. **(C)** t-SNE plot of Utrophin and α-Sarcoglycan features. Each dot is a myotube within a pattern. Shades of green indicates scramble siRNA treatment for non-DMD donors while shades of red indicate scramble siRNA treatment for the DMD donor cells. The blue dots indicate the siRNA3 dystrophin knock down of the non-DMD donor cells. **(D)** Quantification by population Euclidean distance. Euclidean distances were measured using centroid of different populations. **(E)** Using an SVM classifier trained on the scramble siRNA between the DMD and the non-DMD donors, we predicted the siRNA dystrophin KD myotubes. The y-axis shows the prediction of the classifier as non-DMD like myotubes. As evident, by knocking down dystrophin in the non-DMD donor cells, we can generate a phenotype that similar the DMD phenotype (% Like DMD Non-DMD #3 94.8%, Non-DMD #1 – 80.2%).

We used non targeting siRNA (scrambled control) transfected in both DMD and non-DMD donors which were used as controls. We also separately transfected the non-DMD donor cells with anti-Dystrophin siRNA with the purpose of knocking down Dystrophin. The immunofluorescence was performed on Utrophin and α-Sarcoglycan, since this combination gave the best separation quantified by F-Score. As before, we used the features pertaining to Utrophin and α-Sarcoglycan images to determine the effects of Dystrophin knock down.

We quantified the ability of the classifier to accurately classify the different perturbations using F-Score by taking two perturbations at a time (Figure 4B). Treating the non-DMD donor cells with scrambled siRNA did not change the phenotype of these cells as evident from the low F-score values for 10-fold cross validation between scrambled and untreated conditions for the non-DMD donor cells (F-Score 0.8141, Data not shown). We derived a classification model based upon DMD and non-DMD scrambled controls and observed that the combination of Utrophin and α- Sarcoglycan was able to accurately classify the controls with an F-Score of .99 (Figure 4B). Interestingly, a classification model trained on anti-Dystrophin siRNA and the DMD scrambled population was also able to accurately separate the two populations with F-Score of 0.95.

We determined that the cell population treated with the anti-Dystrophin siRNA (Figure 4C, blue dots) is significantly closer in phenotype to the DMD cell population treated with the scrambled siRNA and further away from non-DMD scrambled siRNA phenotype as measured by the Euclidean distance in the feature space and t-SNE plots (Figure 4C-D). This implies that Dystrophin knockdown in non-DMD cells renders a morphological phenotype that is different from non-DMD scrambled phenotype but closer to the DMD-scrambled phenotype.

To further validate that hypothesis that anti-Dystrophin siRNA treatment in non-DMD donors results in a phenotype similar to a DMD patient cell, we developed a classification model between the non-DMD and DMD scrambled controls and used this model to bin the anti-Dystrophin siRNA population (Figure 4E). Very few myotubes from the anti-Dystrophin siRNA were classified into the non-DMD scramble class (5.2% for Non-DMD #3 and 19.8% for Non-DMD #1) indicating that knocking down Dystrophin creates a morphological phenotype that looks like DMD cells.

Taken together, we have demonstrated that morphological profiling in combination with the MyoScreen platform can be used to quantify disease relevant DMD phenotypes.

### Restoration of Dystrophin expression in DMD cells reverses the phenotype associated with the lack of Dystrophin

While knocking down Dystrophin expression in non-DMD cells resulted in a profile movement towards the DMD population, we also tested whether restoration of Dystrophin expression in DMD cells could rescue the observed phenotype.

We used (vivo-Phosphorodiamidate Morpholino Oligonucleomer) vivo-PMO to induce exon skipping in donor cells with permissive mutations as a method of partially restoring dystrophin. Exon skipping is a mechanism that results in restoration of a partial dystrophin in DMD myotubes with certain exon deletions. This PMO induces the spliceosome to ignore a specific exon thereby restoring the reading frame of the Dystrophin gene. We identified DMD donor DMD #1 as the one that is amenable for exon skipping using PMO – exon 45 skipping (Figure 5A).

**Figure 5.**
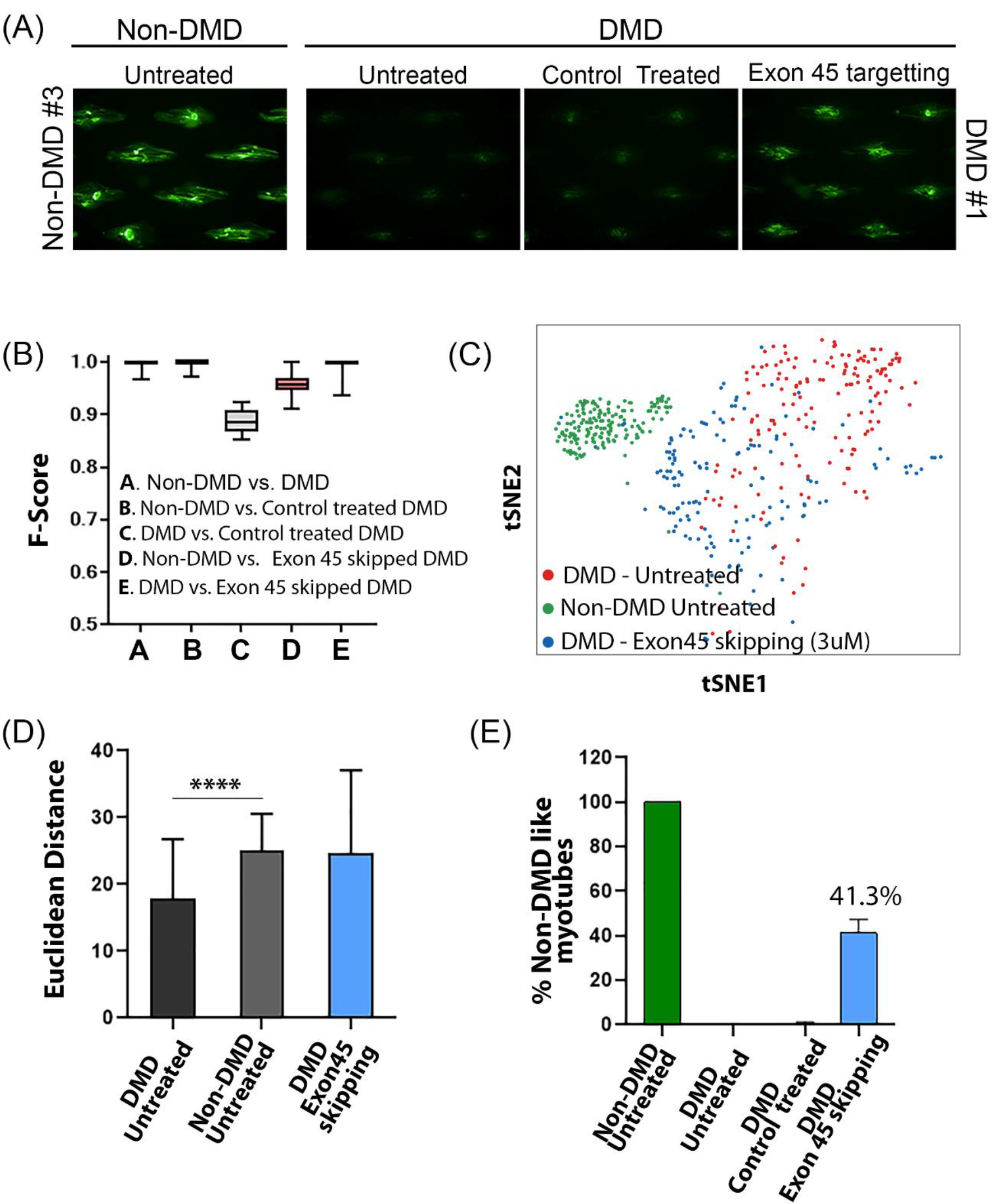
**- Quantification of functional relevance of Dystrophin by exon skipping**. **(A)** DMD donor DMD #1 was chosen for performing exon skipping and restoring partial Dystrophin expression. Images of non-DMD (Non-DMD #3) untreated as well as exon 45 targeting in DMD #1 are displayed. Partial restoration of Dystrophin is observed after exon 45 skipping. All images are pseudo colored. **(B)** F-score from cross validation analysis using SVM linear kernel by taking the 2 different categories at a time using Utrophin and α-Sarcoglycan features. As evident from the F-score, we can separate exon 45 skipped samples from non-DMD controls. **(C)** t-SNE plot of Utrophin and α-Sarcoglycan features. Each dot is a myotube within a pattern. Shades of green indicates non-DMD donor while shades of red indicate untreated DMD donor cells. The blue dots indicate the partially restored dystrophin from DMD #1 donor cells after exon skipping. **(D)** Quantification by population Euclidean distance. Euclidean distances were measured using centroid of different populations. **(E)** Using an SVM classifier trained on the DMD and the non- DMD donors. The y-axis shows the prediction of the classifier as non-DMD like myotubes. As evident, after partial restoration of dystrophin expression in the DMD donor cells, 41.3% of the cells are classified as non-DMD phenotype.

Like our previous analysis, we selected to stain for Utrophin and α-Sarcoglycan and quantified the effects of expressing partial dystrophin using exon skipping. For controls, we introduced a non-targeting PMO in cells from DMD donor DMD #1 (control treated DMD). As shown in Figure 5A, the expression of Dystrophin is negligible in the DMD control cells when compared to non-DMD donor cells. Treatment with the vivo-PMO resulted in restoration of Dystrophin expression (Figure 5A).

We determined that the non-DMD and the non-targeting PMO control treated DMD population displayed a high degree of separation (.99 F-Score), while the DMD and non-targeting PMO control treated DMD showed a lower separation (.88 F-Score) (Figure 5B). Thus, we can infer that a non-targeting PMO does not affect the DMD phenotype.

The DMD and the Exon 45 skipped DMD population displayed a high separation (∼.98 F-score). However, the separation between the Exon 45 skipped DMD and non-DMD population was also high (.95 F-Score). The t-SNE plot in Figure 5C highlights the separation between the three populations – non-targeting PMO-treated DMD, non-DMD and vivo-PMO-treated DMD. As evident from the t-SNE plot, the vivo-PMO-treated DMD cells displayed a different morphological signature when compared to DMD or non-DMD populations. The vivo-PMO treated DMD population shifts towards the non-DMD population, although not completely merging with it. Further evidence of the shift can be obtained from the population mean Euclidean distance to the non-targeting PMO-treated DMD (Figure 5D). This partial shift can be likely due to restoration of a Dystrophin gene sequence that isn’t quite full length, compared to the entire functional dystrophin sequence in the non-DMD population. We built an SVM classifier based on non-DMD and DMD donor population morphological profile and classified individual cells from the Exon 45 skipping DMD population. We determined that 41.3% of the vivo-PMO treated DMD population was classified as non-DMD myotubes (Figure 5E).

Taken together, the results demonstrate the ability of the MyoScreen platform to quantify subtle functional differences between the different DMD populations.

## DISCUSSION

Here, we demonstrate the impact that can be achieved by combining the MyoScreen platform with high-dimensional morphological profiling to enable screening for pathways and targets involved in muscular dystrophy cells with phenotypes established in vitro using physiological systems. Using classical machine learning, we were able to distinguish the DMD and non-DMD populations. Due to the multi-nucleated nature of myofibers, it is difficult to achieve single cell resolution due to challenges in image segmentation. This challenge was overcome using the MyoScreen platform which allowed an analysis of individual islands of cells as one data point, giving us physiological relevance at near single cell resolution.

Previously, several publications have identified Utrophin as a primary target for treating DMD (Loro et al., 2020; Soblechero-Martin et al., 2021). This is based on the finding that Utrophin is upregulated in DMD patients’ skeletal muscle (Kleopa et al., 2006; Taylor et al., 1997). While we observed higher Utrophin expression from one donor (DMD #4) compared to the non-DMD donor cells, this was not found to be consistent across all the DMD donor cells. One possibility for this discrepancy could be due to a biased selection of donors. Our selection process involved donor cells that could be frozen and passaged over time. Despite this discrepancy we also observed Utrophin to be highly relevant in distinguishing the donor populations using morphological profiling. We hypothesize that beyond changes in expression, morphological profiling detected changes in localization of Utrophin within the myofibers between the DMD and non-DMD populations. Beyond Utrophin, we evaluated 6 other proteins in the DAPC and identified Utrophin (Median Cross Validation F-Score: .98) and α-Sarcoglycan (Median Cross Validation F-Score: .92) (Fig 2 and Supplementary S2.2) to be the top two DAPC proteins when trying to distinguish DMD and non-DMD populations, which is in agreement with previous cell genetic models for DMD (Soblechero-Martin et al., 2021). Our method of data analysis not only identified disease relevant proteins/biomarkers but can also rank them based on relevance.

We further used the same method to rank combinations of DAPC proteins as signatures of dystrophic disease. The combination with utrophin for all the different DAPC proteins yielded higher F-scores compared to individual proteins. However, this was not the case with combining DAPC proteins with α-Sarcoglycan, implying less synergy for marker combination with α- Sarcoglycan. One hypothesis for this observation could be due to the location of these proteins in or near sarcolemma (Gawor & Proszynski, 2018) and limitations of microscopy resolution. While the sarcoglycans and dystroglycans are all embedded within the membrane (sarcolemma), Utrophin is localized within the cell near the sarcolemma. As a result, morphological profiling can capture the interaction of Utrophin with sarcoglycans.

We validated our ability to distinguish disease phenotype using two different approaches: (i) By Dystrophin knock down using siRNA in non-DMD donors, and (ii) By introducing Dystrophin in DMD donors by exon skipping (Exon 45).

In the first case, knocking down Dystrophin in non-DMD donor cells yielded a phenotype that is more similar to that of DMD. We observed variations in classification results amongst the donors following siRNA-based knockdown of Dystrophin (94.8% for Non-DMD #3 and 80.2% for Non-DMD #1 DMD like for siRNA dystrophin KD) Figure 4E. This difference can be attributed to donor-to-donor variability even among seemingly “healthy” (non-DMD) donor populations as well as variable efficiency of Dystrophin knock down.

In the second case, we used a DMD donor amenable to exon 45 skipping (DMD #1) and observed that morphological profiling can rescue disease phenotype to a great extent (41.3% as non-DMD for exon 45 skipping treated DMD donor). This result can be explained by the fact that the classification model is binary and requires data to be binned in one of the classes. As shown previously in Figures 5 B-D, the exon 45-skipped population (treated with vivo-PMO) is different from the DMD and the non-DMD population. A binary classification model will not be able to capture this unique population precisely. We can further attribute the observation can be further attributed to the partial restoration of Dystrophin expression (43% dystrophin expression by fluorescence intensity). While the classification percentage is not as high as KD with siRNA, we observe this to be significantly higher compared to the classification of DMD as non-DMD phenotype (0% DMD classified as non-DMD).

Overall, we have demonstrated that the MyoScreen platform together with high dimensional cell profiling enables rapid identification of perturbagens relevant to muscular dystrophy phenotype and could be easily adapted to large scale drug discovery screening.

## MATERIALS AND METHODS

### Reagents

Primary antibody sources are referenced in Supplementary Table S1. Corresponding Alexa Fluor488- or Cy3 or Alexa Fluor647-conjugate polyclonal secondary antibodies were used per supplier’s instructions (Jackson ImmunoResearch, Suffolk, UK & Invitrogen, Courtaboeuf, France). Hoechst 33342 (H3570, Invitrogen) was used to stain nuclei.

### Cell Source and Culture Maintenance

Primary human skeletal muscle myoblasts from Healthy and DMD donors were sourced from different donors (see Table 1) from CBC Biotec Centre de Resources Biologiques, Hospices Civils de Lyon, Lyon, France).

Cells expanded following patient biopsy collection were subsequently enriched for myoblasts using anti-CD56-coated microbeads (Miltenyi Biotec GmbH). Primary vials were sourced, thawed, and the proportion of Desmin postitive cells was determined. Cells were expanded following supplier’s recommendations, maintained in a humidified incubator at 37 °C, 5% CO2, passaged when they reached 70–80% confluence, and cryopreserved into master banks (MB) at which point they were characterized using immunostaining (Desmin positive cells) and the MyoScreen technology (fusion index). Finally, master bank vials were thawed, expanded, and finally cryopreserved into working cell banks (WB). Master banks and working banks cultured in supplier’s recommended conditions were further characterized using immunostaining (Desmin positive cells) and the MyoScreen technology (fusion index). Cells were selected based on consistency in their doubling time, proportion of Desmin positive cells and fusion index.

### High-Throughput Myotube Formation

All steps were accomplished automatically using a Freedom EVO150 workstation (Tecan, Männedorf, Switzerland). Growth medium for Healthy and DMD cells myoblasts was Skeletal Muscle Cell Growth Medium provided by ZENBIO (ZENBIO, Durham, USA).

For all experiments (unless explicitly indicated otherwise), at Day 0, MyoScreen™ plates (CYTOO, France, as described in international patent application WO 2015/091593) containing micropatterns coated with 10 μg/ml fibronectin (Invitrogen) were pre-filled with 100μl/well of growth medium and stored in the incubator at 37°C. Human primary myoblasts were detached from the flasks, counted, and seeded into the plates with 15 000 cells per well in 100μl of growth medium. At Day 1, the growth medium was changed for a differentiation medium well (DMEM/F12 (Invitrogen), 2% horse serum (GE Healthcare, Vélizy-Villacoublay, France), 0.5% penicillin-streptomycin (Invitrogen)), in which myoblasts started differentiating and forming myotubes. Myotubes formation process was then continued for 9 days in differentiation medium without medium replacement.

For experiments performed using standard culture plates, SensoPlates (Dutscher, Brumath, France) were coated with 10 μg/ml fibronectin (Invitrogen) for 2h at room temperature then washed with PBS. Plates were then pre-filled with 100μl/well of growth medium and stored in the incubator at 37°C. Human primary myoblasts were detached from the flasks, counted, and seeded into the plates with 15 000 cells per well in 100μl of growth medium. The day after, growth medium was replaced for a differentiation medium well (DMEM/F12 (Invitrogen), 2% horse serum (GE Healthcare), 0.5% penicillin-streptomycin (Invitrogen), in which myoblasts started differentiating and forming myotubes. Myotubes formation process was continued for 9 days.

### siRNA and vivoPMO treatment and DAPs immunofluorescence staining

After four days of culture, differentiated healthy and DMD myotubes were transfected with either dystrophin and scramble siRNAs (Thermo Fisher Scientific) at 1 nM final concentration using Lipofectamine RNAiMAX (Thermo Fisher Scientific) or with vivo-phosphorodiamidate morpholino oligonucleomers (“vivoPMOs” ; GeneTools - 5’-CCAATGCCATCCTGGAGTTCCTGTAA- 3’) at 3 µM according to respective manufacturers’ instructions.

On day 10, myotubes were fixed for 30 min in 10% formalin (Sigma-Aldrich, St-Quentin-Fallavier, France). Subsequent immunofluorescence staining was according to (Young et al., 2018) PMID 29498891. Myotubes were washed three times in Dulbecco’s Phosphate-Buffered Saline (DPBS, Invitrogen) and permeabilized in 0.5% Triton X-100 (Sigma-Aldrich). After blocking in 1% bovine serum albumin for 20 min (BSA, Sigma-Aldrich), cells were incubated with primary antibodies listed in Table S1 prepared in BSA 1% for 2h at room temperature and then washed three times in DPBS. Secondary antibodies were added for 2 h with Hoechst 33342. Cells were washed three times in DPBS before acquisition.

### Myotube morphology and muscle marker high content analysis

After fixation and immunostaining, quantitative microscopy was performed using the Operetta HCS imaging system with a 10×/0.3 NA objective (PerkinElmer, Courtaboeuf, France). Images were analyzed using scripts developed in Acapella software (PerkinElmer) by the inventors. Eleven fields of view per well were acquired. First, segmentation of myotubes and nuclei were done using respectively the Troponin T or Myosin heavy chain staining and the Hoechst staining.

One to two myotubes per micropattern were usually identified. The threshold of segmentation was set-up to avoid detecting the background noise and eliminate aberrant small myotube structures. At the end of this first step, specific readouts were calculated in the whole well, like the nuclei count and the fusion index (percentage of nuclei included in Troponin T or MHC staining). Usually around 50 to 60 myotubes were detected per well in a control condition. Then, an image clean-up step was performed on the myotubes images to remove myotubes that touch the border of the image. The resulted myotubes were used to extract myotube morphology parameters including the myotube mean area as well as specific marker expression (e.g. dystrophin, utrophin staining intensity).

### Statistical Evaluation

Data are shown as mean ± standard deviation (SD). Prism (GraphPad Software, La Jolla, CA, US) was used for statistical comparisons performed as indicated in figure legends.

**Table S1:** Primary antibodies

| Dystrophin associated protein (DAP) | Antibody Name | Supplier |
| --- | --- | --- |
| α-dystroglycan | Anti-α-Dystroglycan Antibody, clone IIH6C4 | Merck |
| β-dystroglycan | β-dystroglycan, clone 43DAG1/8D5 | Leica |
| α-sarcoglycan | Recombinant Anti-α-sarcoglycan antibody [EPR14773] | Abcam |
| δ-sarcoglycan | δ-sarcoglycan | Leica |
| Syntrophin | SNTB2 Monoclonal Antibody (1351) | Thermo |
| Dysferlin | Recombinant Anti-Dysferlin antibody [JAI-1-49-3] | Abcam |
| Utrophin | Utrophin, clone DRP3/20C5 | Leica |
| Desmin | Recombinant Anti-Desmin antibody [Y66] | Abcam |
| MHC | Myosin 4 Monoclonal Antibody (MF20) | Thermo |
| Troponin T | Polyclonal cardiac troponin T | Abcam |
| Dystrophin | DysB | Leica |

### Cell Profiling analysis

The workflow presented in Figure 2.1 includes the main steps of the machine learning pipeline used in this paper. We start by constructing an annotated database with immunostained plates acquired on a 10x Operetta HCS imaging system. The acquired images undergo a pre-processing step of illumination correction to reduce uneven illumination or background. Then the regions of interest (ROI) are segmented which are in this case, the segmentation of the myotubes dilated by 10 pixels to englobe the membrane. From these ROI are extracted features. The pre- processing, segmentation of region of interest and feature extraction was done in the open-source software CellProfiler (Carpenter et al., 2006). Using machine learning algorithms, we next generate the profile of DMD and non-DMD cells. For machine learning, dimensionality reduction, and data visualization, we used the programming language Python.

### Image Acquisition

The immunostained MyoScreen plates are acquired with a 10x Operetta HCS imaging system. Four channels are acquired: HOECHST 33342 for the nuclei staining, DRAQQ5 for the myotube staining and Cy3 and Alexa 488 channels that are detected to identify DMD biomarkers.

### Image segmentation

The myotube identification is performed through segmentation algorithms on the Troponin T channel. The myotube is considered to be the ROI.

### Feature extraction

For each identified myotube 4 categories of features are extracted that were the most significant: intensity, granularity, texture, and intensity distribution features.

### Intensity features

The first features extracted are the first-order statistics which are calculated from the image histograms. The common features used and proposed are among others: mean, variance, range, intensity on the edge of the ROI, quantile values. These features are extracted from both Cy3 and Alexa 488 channels.

### Granularity features

This set of features is produced by a series of openings of the original image with structuring elements of increasing size. At each step, the volume of the open image is calculated as the sum of all pixels in the ROI. The difference in volume between the successive steps of opening is the granular spectrum. The distribution is normalized to the total volume (integrated intensity) of the ROI.

### Texture features

Texture features measure the degree and nature of textures within the myotubes to quantify their roughness and smoothness. This set of features measures information regarding the spatial distribution of the various channel intensity levels. A region of interest without much texture has a smooth appearance; a region of interest with a lot of texture will appear rough and show a wide variety of pixel intensities. To calculate this set of features we used Haralick texture features (Haralick et al., 1973)that are derived from the co-occurrence matrix. The matrix contains information about the correlation of intensity between one pixel and the one placed n-pixel further.

### Intensity distribution features

This feature set measures the spatial distribution of intensities within each object. Given an image with identified ROI, this set of features measures the intensity distribution from each object’s center to its boundary within a set of rings. The distribution is measured from the center of the object, where the center is defined as the point farthest from any edge.

### Profile generation

The generated datasets are standardized and balanced to have approximately the same number of ROI in non-DMD and DMD cells categories. The datasets used for the training of the model are composed solely of the control conditions and contain the set of detected myotubes with its extracted features.

The evaluation metric used to predict model’s efficacy to predict the category of a myotube is the F-score. The F-score is a measure that evaluates the model’s accuracy and is defined as: 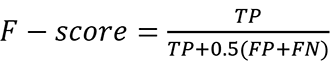 where TP = True Positive, FN = False Negative and FP = False Positive values.

A F-score of 1 depicts a perfect classification: the two categories, non-DMD and DMD, are completely distinguishable. A F-score below 0.9 is considered to be low and not allowing a good separation between the non-DMD and DMD phenotypes.

The evaluation of the model is performed 10 times over the dataset which is randomly split each time in 90% used for the training of the model and 10% that is used to test the model’s accuracy.

### Machine learning algorithms

We chose the Support Vector Machine (SVM) algorithm to distinguish myotubes coming from non-DMD and DMD categories. In the SVM algorithm (Cortes & Vapnik, 1995), each data item is plotted in the n-dimensional space of the features (n being the number of features extracted for each segmented ROI) and it aims at obtaining a hyperplane that will optimally separate the items from the two non-DMD and DMD categories.

### % Of non-DMD like myotubes readout definition

The % of non-DMD like myotubes is defined as the percentage of myotubes out of the total that are predicted to have a non-DMD phenotype by the model. The predictions are generated on the datasets that do not belong to the training set and the output for each myotube is a probability between 0 and 1 of belonging to one class or the other. If the probability for one myotube is below 0.5, we consider that this myotube belongs to one category while if it is higher or equal than 0.5 we consider that it belongs to the other category.

### Data visualization

The representation of a myotube features dataset is done using the t-distributed stochastic neighbor embedding (t-SNE) dimensionality reduction method (van der Maaten & Hinton, 2008). The data in Figures 3, 4, 5 is represented using the t-SNE method for visualization of high-dimensional features in a 2D space.

## ACKNOWLEDGMENTS

We thank the Centre de Ressources Biologiques CBC Biotec, Hospices Civils de Lyon, Lyon, France, for providing the primary human skeletal muscle myoblasts.

---

**Figure S2.1.**
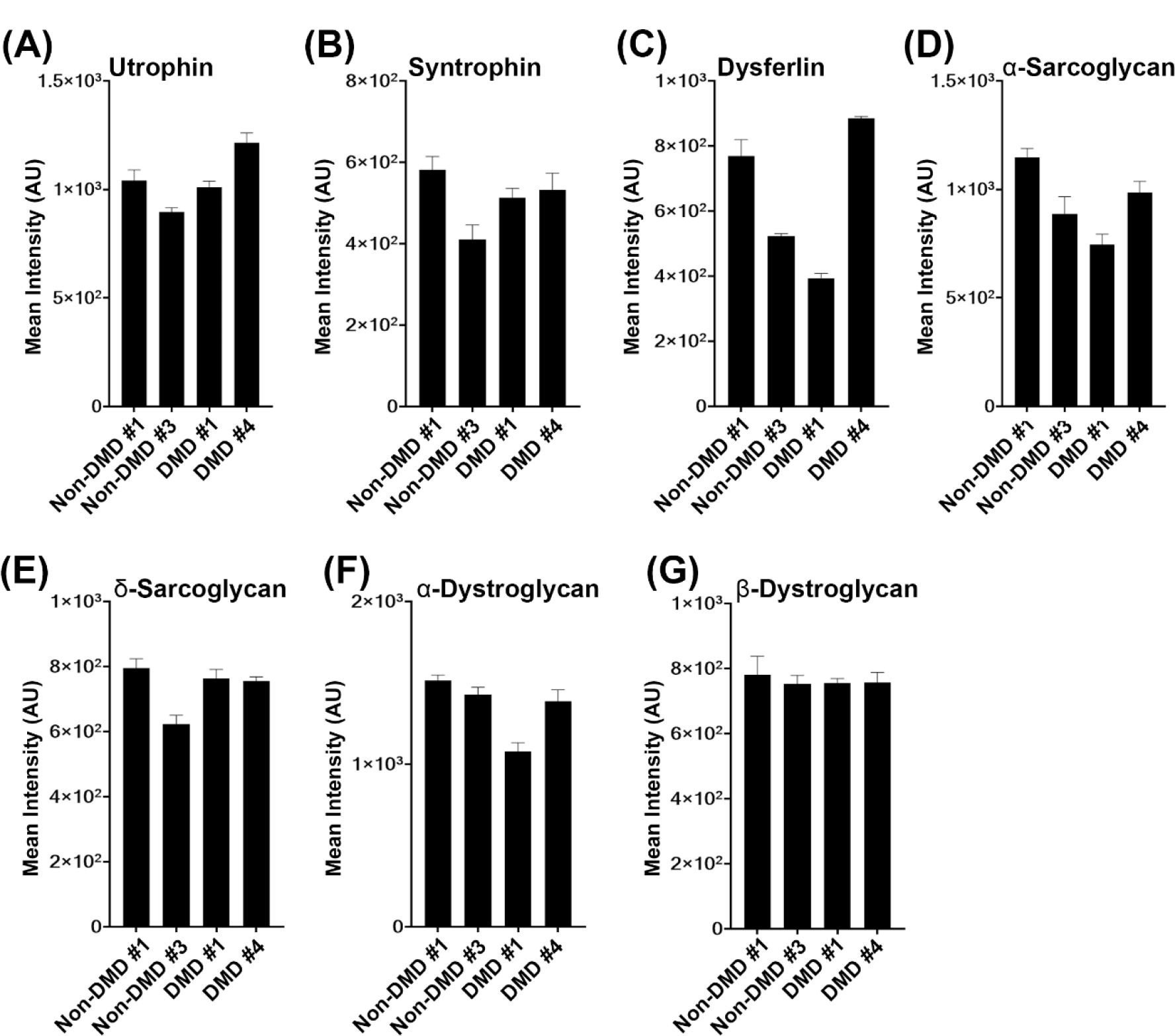
**Quantification of DAPC protein expression across different donors**. **(A-G)** Average pixel intensity of individual DAPC protein (a measure of DAPC protein expression level) across multiple myotubes of each DAPC protein. Significant differences in signal intensity were not detected for any of the DAPC proteins analyzed.

**Figure S2.2.**
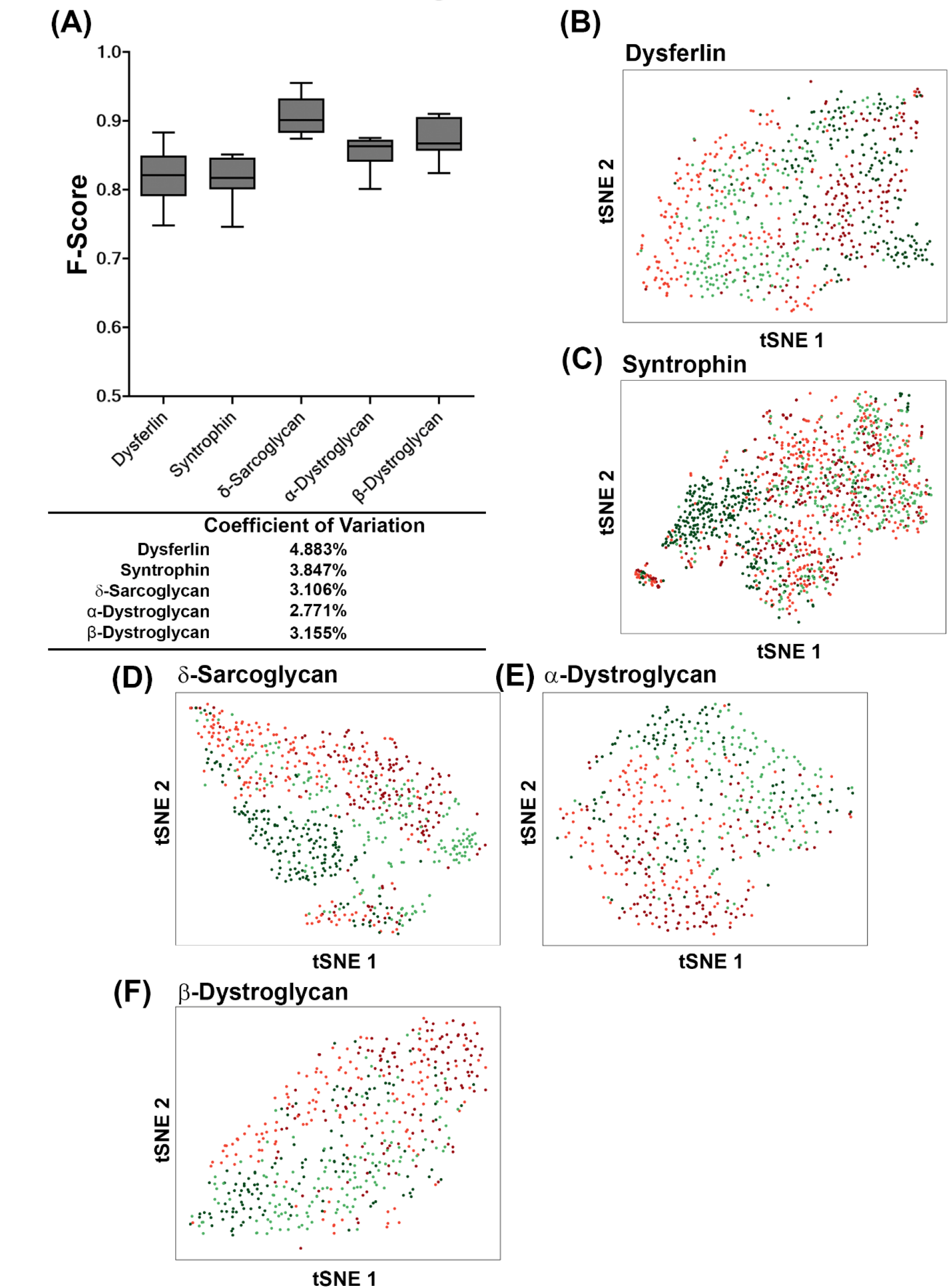
**Cross validation accuracy for other DPAC proteins excluding Utrophin and a- Sarcoglycan**. (A) Cross validation using the subcellular features using individual DAPC proteins obtained from cell profiling from cells cultured on MyoScreen platform. The corresponding t-SNE plates are displayed in the different plots.

**Figure S2.3.**
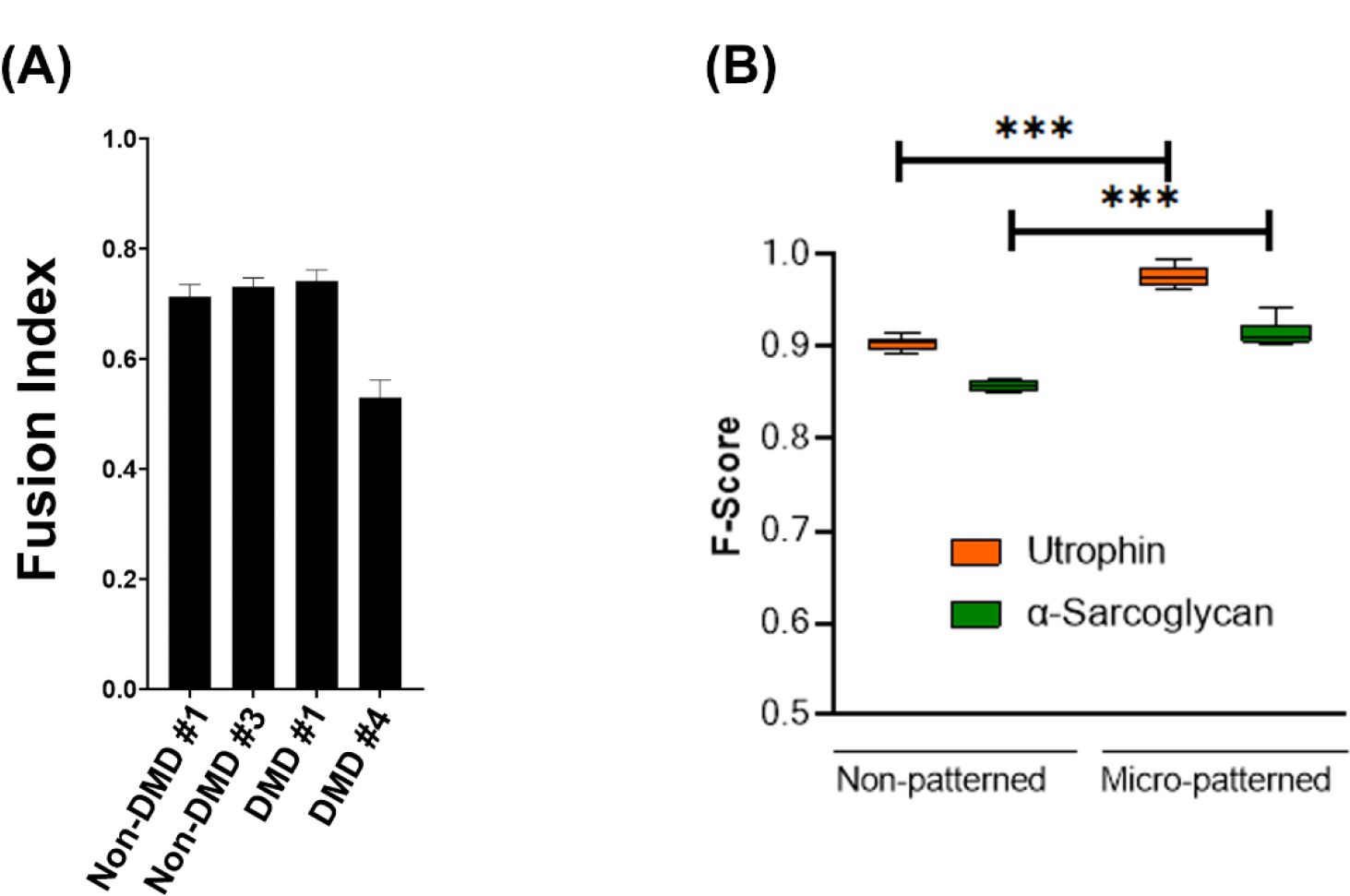
**Validation of the MyoScreen profiling platform**. **(A)** Fusion index of myotubes cultured using standard 96 well plates. No significant differences between DMD and non-DMD donors were observed. **(B)** Cross validation using the subcellular features obtained from cell profiling from cells cultured in standard 96 well plates compared with cells cultured in the MyoScreen platform. A significantly higher cross validation F-Score can be observed when cells are cultured in the MyoScreen platform.

